# DNA-binding and dimerization of the SOG1 NAC domain are functionally linked with its ability to undergo liquid-liquid phase separation

**DOI:** 10.1101/2025.05.09.653017

**Authors:** Kim Mignon, Rani Van der Eecken, Margot Galle, Manon Demulder, Joris Van Lindt, Lieven De Veylder, Henri De Greve, Remy Loris

## Abstract

Liquid-liquid phase separation is a key phenomenon in the regulation of transcription in eukaryotes leading to the formation of so-called membraneless organelles. While transcription factors take part in several types of membrane-less organelles, it remains unclear how specific DNA binding, multivalent interactions with DNA/RNA and condensation are interlinked. Here we show that the NAC domain of SOG1 (SOG1^NAC^), a transcription factor that is central to the DNA damage response in plants, can undergo liquid-liquid phase separation *in vitro* in the presence of both RNA or double stranded DNA. This behaviour, as well as the ability of SOG1^NAC^ to bind DNA in a sequence-specific manner are dependent on its potential to form homodimers and the presence of a cluster of positive charges in its DNA binding site. Short double-stranded DNA fragments containing the sequence motif that is specifically recognized by SOG1^NAC^ inhibit RNA-mediated phase separation, suggesting overlapping binding sites for DNA and RNA. This may reflect a complex interplay between DNA and RNA binding that could control the formation of condensates at transcription sites.

## Introduction

Many cellular functions such as RNA metabolism and processing, ribosome biosynthesis, gene regulation and stress responses involve the formation of membrane-less organelles (MLOs) ((1), (2), (3), (4), (5)). The latter are biomolecular condensates of specific macromolecular components that are formed via a process of liquid-liquid phase separation (LLPS). MLOs are typically dynamic in composition with fast exchange of components with their surroundings. The entropic cost associated with the formation of MLOs is overcome by favourable multivalent interactions between the biomolecules they consist of (1), (6).

Membraneless organelles are implicated in transcriptional regulation during which many macromolecules, including transcription factors and RNA species cluster at specific genomic locations (for a review see (7)). In general, eukaryotic transcription factors (eTFs) recognize and bind specific DNA motifs of 5-15 base pairs (bp) in promoter or enhancer sequences with their folded DNA-binding domains (8). Other components of the transcription machinery like coactivators, regulatory cofactors, and RNA polymerase II are recruited by multivalent protein-protein interactions with the intrinsically disordered transactivation domains (TAD) of eFTs (9, 10). Henninger *et al.* (2021) proposed that the formation of transcription condensates is controlled by an RNA-mediated feedback mechanism. Low concentrations of RNA produced during transcription initiation promote condensation, while the burst production of mRNA during elongation leads to dissolution of the condensates (8, 10, 11). Research on LLPS of transcription factors has mostly focused on their intrinsically disordered domains. The latter often harbour low complexity regions that create multivalency and are more than folded domains prone to post-translational modifications. Both these properties haven proven to be key in LLPS (for reviews see (12, 13)). Also folded domains, and especially RNA and DNA binding domains of LLPS-prone proteins have been observed to influence the formation of MLOs (14–16). However, the link between LLPS and the ability of folded nucleic acid binding domains to specifically or non-specifically bind DNA or RNA is poorly understood.

SUPPRESSOR of GAMMA RESPONSE 1 (SOG1) belongs to a large plant-specific family of NAC [NO APICAL MERISTEM (NAM), *ARABIDOPSIS* TRANSCRIPTION ACTIVATION FACTOR (ATAF), CUP-SHAPED COTYLEDON (CUC)] transcription factors that function in growth and development or respond to abiotic and biotic stresses (17, 18). Among these, SOG1 activates more than 300 genes involved in DNA repair, cell cycle arrest or apoptosis in response to DNA damage in the model plant *Arabidopsis thaliana* (19, 20). The protein consists of a conserved, folded DNA binding domain of 155 amino acids (called the NAC domain) followed by a highly divergent C-terminal domain (CTD) that is intrinsically disordered (21, 22). SOG1 contains an additional 57 amino acid extension of unknown function that is lacking in most other NAC transcription factors (19, 23). The transcriptional activation function of SOG1 is controlled by the phosphorylation of five SQ sites by the DNA damage sensing kinases ATM and ATR and by phosphorylation of a single CASEIN KINASE 2-dependent site (24–27).

In this study we investigate the *in vitro* DNA binding of the folded NAC domain of *Arabidopsis thaliana* SOG1 (which will be called SOG1^NAC^ from now on). Next, we link its specific double-stranded DNA (dsDNA) binding to its potential to form functional homodimers and its ability to undergo DNA- and RNA-driven LLPS. In summary, we investigate the putative correlation between SOG1^NAC^ dimerization, specific and non-specific DNA/RNA binding and the potential for phase separating behaviour. We conclude that the NAC domain of SOG1 undergoes LLPS in the presence of RNA and dsDNA, and that this potential requires an intact DNA binding site and the formation of dimers.

## Materials and methods

### Proteins and nucleic acids

The SOG1 amino acid (AA) sequence used in this paper corresponds to UniProt accension code Q6NQK2. The SOG1^NAC^ domain ranges from L58 to Q212. An overview of the DNA sequences that were used in the various experiments of this paper can be found in Table 1. The single-stranded DNA (ssDNA) oligos of these sequences (and their reverse complements) and the used mRNA proxy called Polyadenylic acid potassium salt (poly-A, CAS number: 27416-86-0) were obtained from Sigma-Aldrich.

**Table 1:**
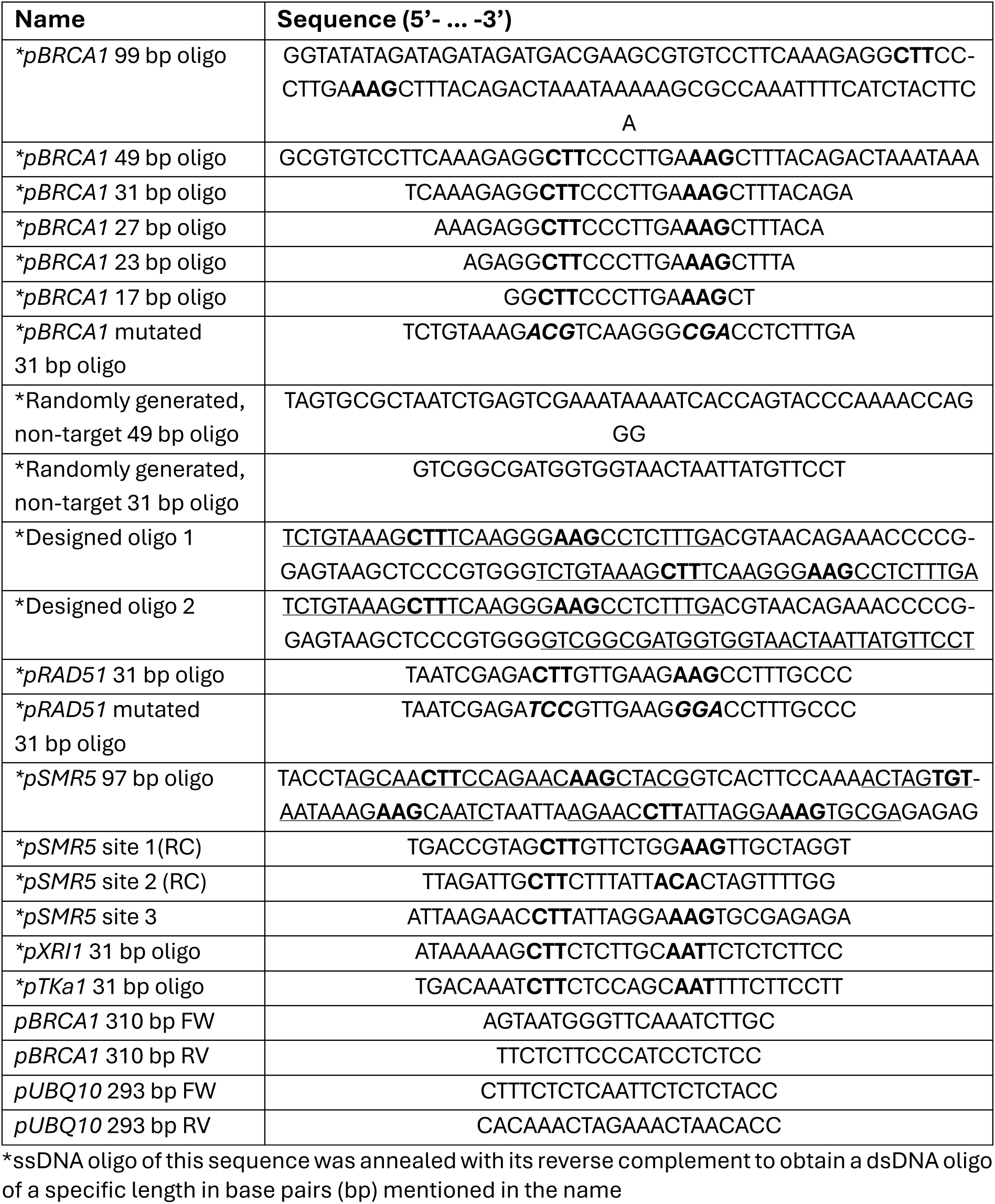
Sequences of the DNA oligos used in this paper. The six CTT-AAG nucleotides of the CTT(N7)AAG “Ogita motif” or variations of these nucleotides are highlighted in bold text. Mutated nucleotides are written in italic. The reoccurring 31 bp oligos present in the two “Designed oligos” are underlined. The three 23bp potential binding sites within the pSMR5 97 bp oligo are also underlined. RC = reverse complement, to represent the binding motifs more clearly.

### Structure prediction of the SOG1^NAC^-dsDNA complex

The structure of SOG1^NAC^ in complex with the *pBRCA1* 31 bp oligo was predicted using AlphaFold3 (28). For this, two copies of the SOG1^NAC^ protein as well as one copy of the *pBRCA1* 31 bp ds-oligo were entered in https://alphafoldserver.com/. Visualization and surface charge mapping were performed using PyMOL and the APBS Electrostatics plugin delivered by PyMOL (29).

### Cloning of the SOG1^NAC^ construct and its mutants

The gene sequence of SOG1^NAC^ preceded by a 6xHis-tag was synthetically produced and cloned into the pET-11d vector via restriction-ligation (NcoI and BamHI) by Genscript. Site-specific mutagenesis was performed on this clone to obtain the three SOG1^NAC^ mutant clones: SOG1^NACΔ1-6^, SOG1^NAC-RKRR^ and SOG1^NAC-DNAstrand^. In contrast to wild type SOG1^NAC^, the SOG1^NACΔ1-6^ mutant protein lacks the first six amino acids (ΔL58-K63). The SOG1^NAC-RKRR^ mutant contains following four mutations: R79A, K80A, R81A, and R82A. The SOG1^NAC-DNAstrand^ mutant harbours following five mutations: R93S, H95S, T97A, R99S and T100S. To obtain these three mutant clones, specific mutant primer sets were designed (Supplementary Table 1). The mutated reverse primers were used in combination with a forward primer containing an XbaI restrictive site (“*XbaI FW”*) to amplify the sequence of the SOG1^NAC^ clone upstream of the mutations with homologous regions of the pET-11d vector. The mutated forward primer was used in combination with a reverse primer with a mutated BamHI restrictive site to Acc651 (“*BamHI (to Acc651 mutated) RV”)* to amplify the sequence of the SOG1^NAC^ clone downstream of the mutations with homologous regions of the pET11d vector. Via overlap PCR using primers *XbaI FW* and *BamHI (to Acc651 mutated) RV*, the whole coding sequences of the mutants with a 6xHis-tag were obtained. Subsequently the construct was cloned into the pET-11d vector at the XbaI and BamH1 site via Gibson Assembly. Positive colonies were selected with restriction enzyme Acc651 and correct insertion was confirmed by Sanger sequencing.

### Expression and purification of the SOG1^NAC^ domain and its mutants

*E. coli* BL21(DE3) cells, transformed with a pET-11d vector carrying the gene sequence of SOG1^NAC^ or its mutants with an N-terminal 6xHis-tag, were inoculated in 1L LB medium with ampicillin (100 µg/ml) and grown at 37 °C until an OD_600 of_ 0.6-0.8 was reached. Protein expression was induced by adding 0.5 mM isopropyl β-D-1-thiogalactopyranoside (IPTG) and the cultures were incubated at a temperature of 16 °C for wild type (WT) SOG1^NAC^ and SOG1^NACΔ1-6^ mutant and 23 °C for the SOG1^NAC-RKRR^ and SOG1^NAC-DNAstrand^ mutants. Cells were collected by centrifugation (5000 x g, 30 min) the next morning, except for the SOG1^NAC-DNAstrand^ mutant for which 2 hours of incubation was sufficient for optimal expression. Pellets of 2 L bacterial culture were resuspended in 30 mL lysis buffer (50 mM Tris pH 7.5, 500 mM NaCl, 10 % Glycerol, 2 mM Imidazole, 2 mM beta ME) supplemented with ½ protease inhibitor tablet (EDTA free cOmplete™ ULTRA tablets, Roche). Resuspended pellets were used immediately for purification or were flash frozen and stored at −80 °C until use.

After thawing, the pellets were incubated for 20 min rotating at room temperature after adding 2 mM MgCl_2 an_d 50 µg/ml DNase. Hereafter, the cells were lysed by sonication for 5 min with 5 sec ON/5 sec OFF cycles at 80 % amplitude using a VCX-70 Vibra Cell (Sonics). Next, samples were centrifuged at 35 000 x g for 45 min at 4 °C. The supernatant was filtered (0.45 µm) and loaded on a 1 mL HisTrap™ HP column (Cytiva) that was equilibrated prior with the lysis buffer. After loading the sample, the column was washed with lysis buffer. When the UV_280 st_abilized, a step gradient over 6 column volumes (CV) of 2 %, 6 CV of 50 % and 6 CV of 100 % elution buffer (50 mM Tris pH 7.5, 500 mM NaCl, 10 % Glycerol, 1 M Imidazole, 2 mM beta ME) was applied to elute the bound proteins. The eluted fractions containing the protein of interest (POI) were loaded on a Superdex 75 16/90 column (GE Healthcare) equilibrated with gel filtration buffer (20 mM NaPi pH7.5; 150 mM NaCl; 1 mM TCEP). Peak fractions of the POI were pooled, flash frozen and stored at −80 °C.

### Circular dichroism (CD)

Protein samples were buffer exchanged to 10 mM NaPi, pH 7.5 by dialysis. CD spectroscopy of the POI samples was performed on a Biologic MOS-500 spectropolarimeter using a 1 mm quartz cuvette (Hellma) at a POI concentration of approximately 0.25 µg/µl. Spectra were collected at 25 °C and at wavelengths ranging from 190 nm to 260 nm in steps of 1 nm. The buffer spectrum was subtracted from the raw CD data (both θ in mdeg) and the resulting CD spectra were normalized to molar ellipticity ([θ] in deg cm² dmol-1 res-1) using following formula 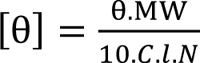: with MW the molecular weight in Da, C the concentration in mg/ml, l the cuvette path length in cm and N the number of amino acids.

### Analytical size-exclusion chromatography (SEC)

Analytical SEC was performed on a Superdex 75 Increase 10/300 column (GE Healthcare) equilibrated with 20 mM NaPi pH 7.5, 150 mM NaCl and 1 mM TCEP. For each POI, the column was loaded with 500 µl of protein at a concentration of approximately 2 mg/ml. In a separate run, 200 µl of the Bio-Rad Gel Filtration Standard was loaded under the same conditions as those of the protein samples under investigation. Based on the elution volumes of the proteins present in the standard, the apparent molecular weights (MW) of the proteins in solution were determined. The apparent MW has the following linear relationship with the Kav retention factor (30):

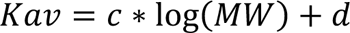

The latter is determined by following formula 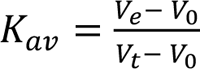: with V_e th_e elution volume, V_0 th_e void volume and V_t th_e total column volume. For each SEC set-up, the exact linear relationship is defined by plotting the logarithm of the MW values of the proteins in the standard (bovine thyroglobulin = 670 kDa, bovine γ-globulin = 158 kDa, chicken ovalbumin = 44 kDa, horse myoglobin = 17 kDa and vitamin B12 = 1.35 kDa) as a function of its Kav-value and fitting a linear regression curve (using the least square method) through these data points. Finally, using the equation of the regression curve, the apparent molecular weights of the proteins in solution could be determined based on their elution volumes.

### Dynamic light scattering (DLS)

DLS measurements were performed using a DynaPro NanoStar (Wyatt) device and the data were analysed using the DYNAMICS 7.10.1.21 software. Growth of droplets was followed by adding poly-A to spin-filtered SOG1^NAC^ to induce LLPS. A reaction with following reaction conditions was set up in triplicate: 20 µM protein and 25 µg/ml poly-A in 9.1 mM phosphate buffer, 68 mM NaCl, 0.45 mM TCEP. Every 5 minutes, five acquisitions of 8 seconds were measured for 4 hours at room temperature.

### Electrophoretic mobility shift assay (EMSA): regular and ^32^P-labelled

Unlabelled dsDNA fragments used during EMSA experiments were obtained by hybridising two complementary ssDNA oligos for 10-15 min at 99 °C and subsequently letting the sample cool down slowly to room temperature (duration approximately 1.5 h). Varying concentrations of POI and hybridised dsDNA at 2 µM were mixed in 20 mM sodium phosphate pH 7.5, 150 mM NaCl and 1 mM TCEP. Binding reactions were carried out in a total volume of 10 µl at 25 °C for 25 min. Hereafter, 2 µl of loading dye (25 % ficoll, 0.1 % xylenexyanol and 0.1 % bromophenol) was added to each sample. Samples were run on a 13 % native polyacrylamide gel prepared with 1 x TBE (89 mM Tris Base, 89 mM Boric acid, 2.5 mM EDTA) that was pre-ran at 100 V for 30 min. Electrophoresis was performed at 120 V until the fastest migrating dye reached 2/3^rd^ of the length of the gel. Gels were stained for 15 min with ethidium bromide or GelRed® for DNA visualisation.

For radioactive EMSAs with SOG1^NAC^, a (5’-^32^P) single-end-labelled 99 bp DNA fragment of *pBRCA1* was used. The ssDNA oligo of this fragment was first labelled with [γ ^32^P]-ATP (PerkinElmer) by T4 polynucleotide kinase (Fermentas). Next, dsDNA was obtained by hybridizing the labelled ssDNA oligo with the complementary oligo using the method described previously. Subsequently, the hybridized, labelled, dsDNA fragment was purified from gel (6 % polyacrylamide gel electrophoresis) as described in (31). Varying concentrations of SOG1^NAC^ and labelled DNA at 15 000 counts per minute (cpm) were mixed in 20 mM sodium phosphate pH 7.5, 1 M TCEP and 50, 150, 300 or 500 mM NaCl. Binding reactions were carried out in a total volume of 20 µl at 25 °C for 25 min. After the addition of 3 µl loading dye (25 % ficoll, 0.1 % xylenexyanol and 0.1 % bromophenol), samples were run on a 10 % polyacrylamide gel prepared with 1 x TBE (89 mM Tris Base, 89 mM Boric acid, 2.5 mM EDTA) that was pre-ran 100 V for 30 min. Electrophoresis was performed at 120 V until the fastest migrating dye reached the bottom of the gel. X-ray sensitive films were used for visualisation.

### Isothermal titration calorimetry (ITC)

ITC measurements were carried out on a MicroCal PEAQ-ITC system (Malvern) at 25 °C and in 20 mM NaPi pH 7.5, 150 mM NaCl and 1 mM TCEP. The DNA samples (obtained in the same way as for the unlabelled EMSA experiments) were loaded into the sample cell at concentrations ranging between 6 and 9 µM. Protein samples were loaded into the syringe at concentration ranging between 155 and 190 µM. For the measurements, a reference power of 10 µcal/s and a stirring speed of 750 rpm were used. Each measurement entailed 20 injections of 2 µl with 150 s intervals, preceded by a first test injection of 0.4 µl. Prior to the start of each run, an initial delay of 300 s was included. For data integration and determining the best fitted binding model for the resulting binding isotherms, the MicroCal Peaq-ITC Analysis Software (version 1.41, Malvern Panalytical) was used. In all cases the “one set of sites” model was used for fitting the binding isotherms, which assumes the presence of one binding site (protein dimer/dsDNA) or any number of sites with the same K_D an_d ΔH. The equations describing this model are provided in the Malvern Microcal PEAQ-ITC user manual (32).

### Turbidity assay

Absorbance of mixed protein and poly-A samples in total volume of 30 µl was measured in triplicate at 390 and 600 nm in non-binding 384 black well plates with transparent bottom (Greiner bio-one, µClear®). Measurements were performed on a SynergyTM Mx plate reader at room temperature in 9.1 mM phosphate buffer, 68 mM NaCl, 0.45 mM TCEP. Phase separation was induced by addition of poly-A right before starting the measurement for minimum 30 minutes.

### Microscopy

Droplet formation was visualized using a Leica DMi8 microscope connected to a Leica DFC7000 GT camera. Reactions of 25 µl in volume (25 mM phosphate, pH 7.5, 75 mM NaCl, 0.5 mM TCEP or 9.1 mM phosphate, pH 7.5, 68 mM NaCl, 0.45 mM TCEP) with fluorescently labelled proteins and/or nucleic acids and unlabelled molecules in excess were prepared in non-binding 384 black well plates with transparent bottom (Greiner bio-one, µClear®). Prior to visualization with an 100x immersion oil objective, the plates were incubated on ice for 10 minutes. The SOG1^NAC^ protein and its mutants were labelled with Dylight 488 (Thermofisher scientific™) according to the manufacturer’s protocol. Poly-A was labelled with Cy5 using pCp-Cy5 (Jena Bioscience) and T4 RNA ligase (Thermofisher scientific™). Cy5-labeled DNA oligos were obtained from Integrated DNA Technologies. Green fluorescence was detected by using a FITC filter. Red fluorescence was detected by a Rhodamine filter. All experiments were performed in duplicate.

### Fluorescence recovery after photobleaching (FRAP)

Samples preparation is identical to the one described above for the microscopy experiments. Firstly, 10 pre-bleaching images were taken every 50 msec. Next bleaching was performed for 50 msec with a laser at 488 nm with 100% intensity. Finally, post-bleaching images were recorded each second for 2-3 minutes.

### Biolayer interferometry (BLI)

BLI competition assays using nucleic acids, DNA and RNA, and POI bait were performed using an Octet 96 Red instrument in 10 mM phosphate, pH 7.5, 75 mM NaCl, 0.5 mM TCEP at room temperature while shaking at 1000 rpm. Ni-NTA biosensors (Satorius) were loaded with 5 µg/ml of 6xHis-tagged SOG1^NAC^ or its mutants to attain a shift between 4 and 4.5 nm, followed by a baseline in buffer for 120 seconds. Next, association of 0.5 µM poly-A for 240 seconds and a second association with different concentrations BRCA1 or random 31 non-target oligo was performed.

## Results

### The SOG1^NAC^ dimer binds specifically to DNA *in vitro*

The *in vivo* DNA binding specificity of *Arabidopsis thaliana* SOG1 has previously been investigated (19, 20). Using ChiP-seq and microarray experiments, Ogita et al. (2018) proposed the palindromic consensus sequence CTT(N)_7AA_G as the binding motif of SOG1 (19). This “Ogita motif”, however, occurs only in 51 % of the identified SOG1 targets. In a different study, Bourbousse et al. (2018) identified enriched sequence motifs present in the promoters of gene groups that are upregulated by SOG1 in response to genotoxic stress, as potential SOG1 binding sites (Supplementary Figure S1) (20). The “Bourbousse motifs” are similar to each other. And despite being less restrictive than the CTT(N)_7AA_G consensus sequence, the latter is nevertheless included in all three of them, confirming this sequence as a putative binding motif for SOG1.

To confirm these *in vivo* results *in vitro*, EMSA experiments were carried out using SOG1^NAC^ and a 49 bp, ds fragment of the *BRCA1* gene promoter (*pBRCA1*), an established *in planta* target of SOG1 (Figure 1A, Table 1) (19). The *pBRCA1* fragment contains the putative CTT(N)_7AA_G Ogita motif and matches with the Bourbousse motifs (Supplementary Figure S1). SOG1^NAC^ shows specific binding to the *pBRCA1* fragment, but not to a randomly generated dsDNA fragment of the same length lacking the Ogita and Bourbousse motifs (Table 1) - from now called non-target DNA (Figure 1A). For the latter, only a transient protein-DNA complex is visible, which is likely formed via non-specific electrostatic interactions between the negatively charged DNA and the overall positively charged NAC domain(Figure 2D). This complex is also visible with *pBRCA1* DNA in presence of excess SOG1^NAC^ (Figure 1A).

**Figure 1:**
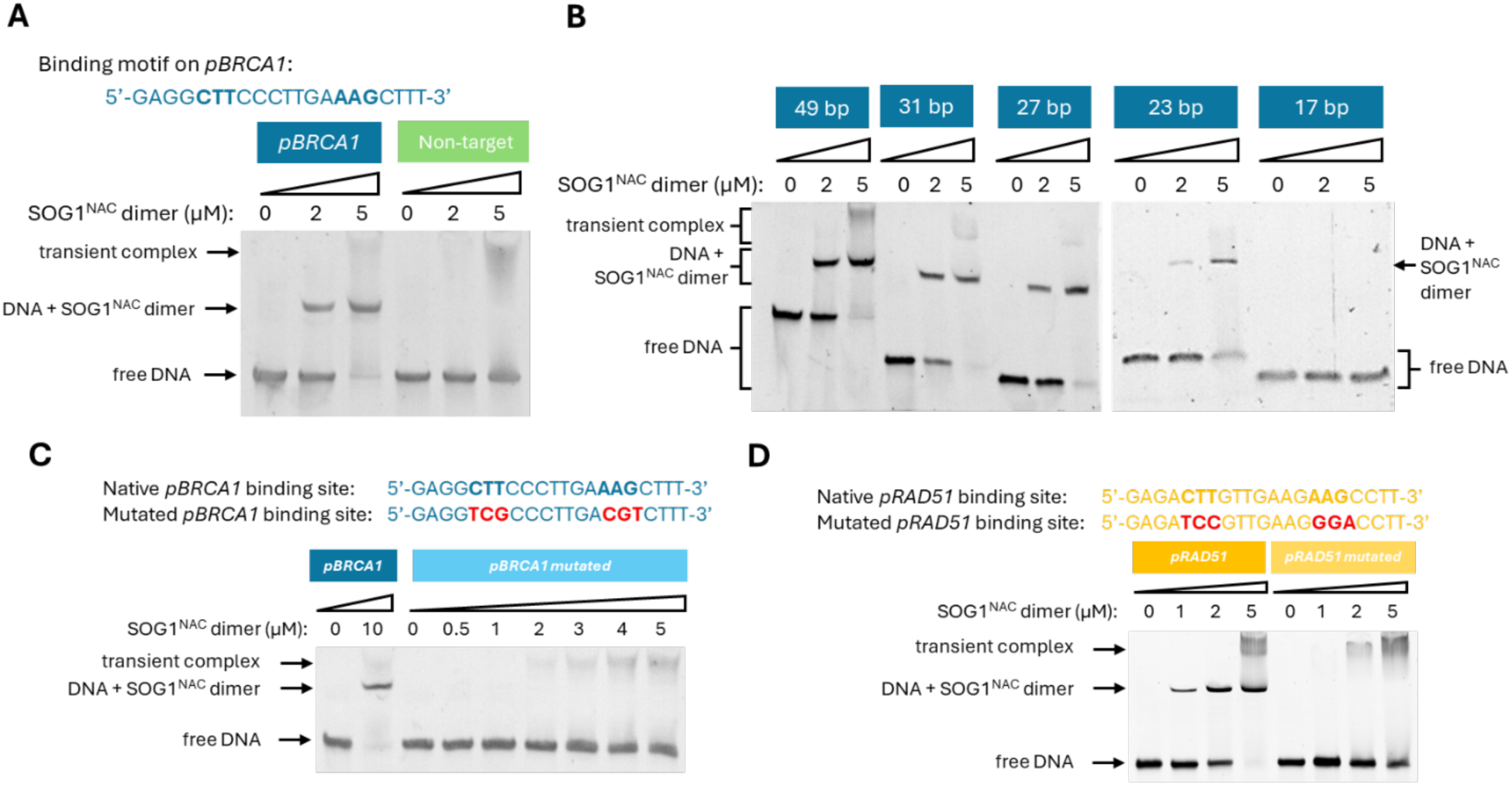
SOG1^NAC^ binds specifically to DNA (2μM) in vitro. (A) Binding motif located on pBRCA1 and EMSA of SOG1^NAC^ with a 49 bp dsDNA fragment of pBRCA1 DNA with this motif and a 50 bp fragment of randomly generated, non-target dsDNA. (B) EMSA of SOG1^NAC^ with pBRCA1 dsDNA oligos of decreasing length, all centrally containing the binding motif. (C) EMSAs of SOG1^NAC^ with a WT and mutated 31 bp ds oligo of pBRCA1. (D) EMSAs of SOG1^NAC^ with a WT and mutated 31 bp ds oligo of pRAD51.

**Figure 2:**
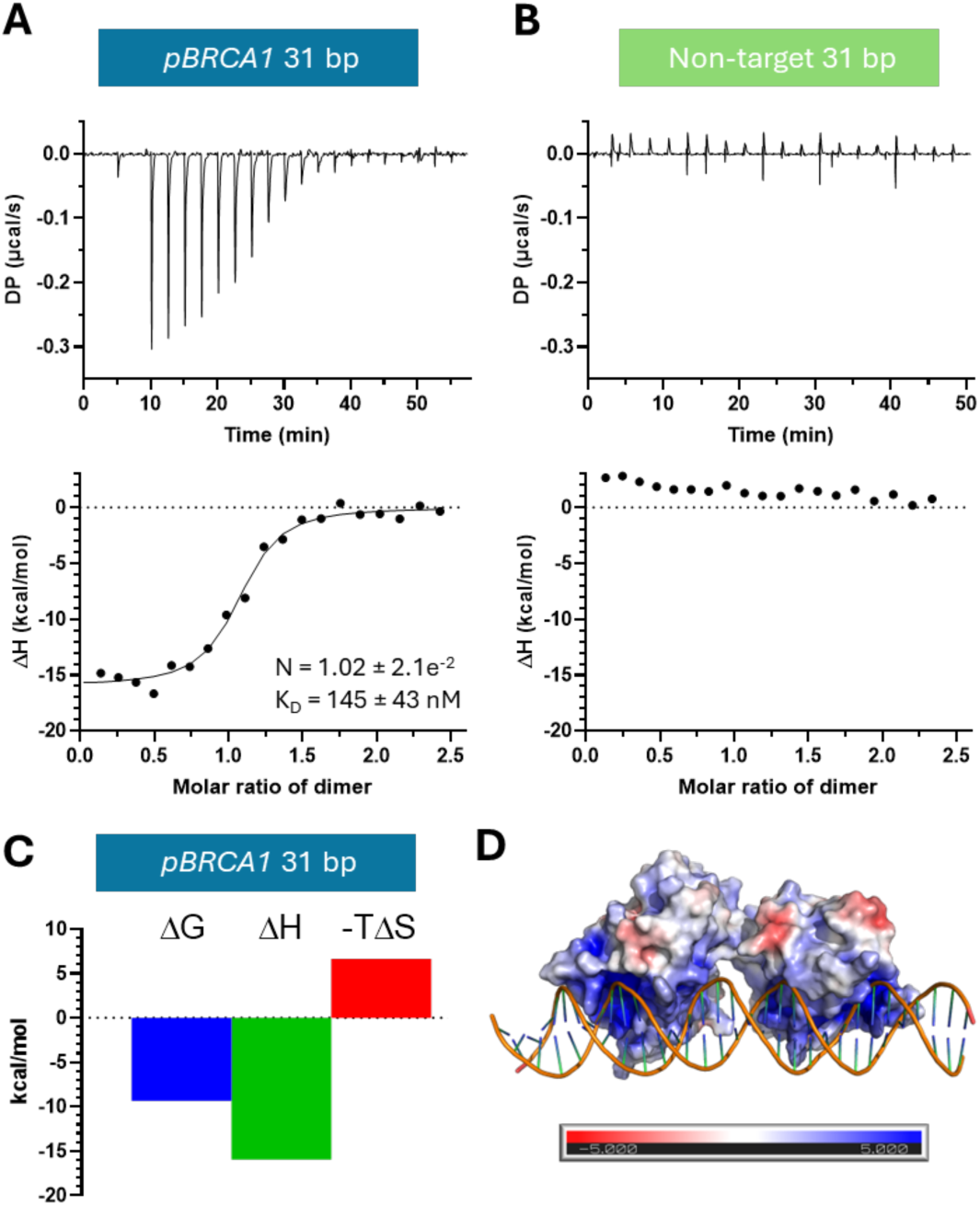
The SOG1^NAC^ dimer binds with nanomolar aZinity to pBRCA1 DNA. (A) & (B) ITC of SOG1^NAC^ with a 31 bp pBRCA1 ds oligo (containing the complying binding site) and a randomly generated, non-target dsDNA fragment of 31 bp, respectively. Upper graphs show the raw heat plot, lower graph show the integrated heat plot. (C) Signature plot of SOG1^NAC^ binding to target 31 bp pBRCA1 DNA. (D) Surface charge map of the SOG1^NAC^ dimer in complex with dsDNA, as predicted by AlphaFold3.

The specificity of SOG1^NAC^ for the binding site on the *pBRCA1* fragment was further confirmed by probing two designed 95 bp dsDNA oligos: one containing two 31 bp *pBRCA1* binding sites connected by a random linker and one containing only one such 31 bp site connected by the same linker to a 31 bp non-target DNA fragment (Table 1, Supplementary Figure S2). The data obtained indicates that SOG1^NAC^ specifically targets the binding site on the *pBRAC1* fragment as two bands of specifically bound protein-DNA complex are present for the first DNA oligo, whereas only one band is observed with the second oligo.

The binding site of SOG1^NAC^ on *pBRCA1* was further delineated using fragments of decreasing lengths. The minimal length of the SOG1^NAC^ binding site lays between 17 and 23 bp (Figure 1B, Table 1). The six CTT-AAG nucleotides that are separated by the N_7-l_inker in the CTT(N)_7AA_G Ogita motif appear to be critical for binding of *pBRCA1* as mutations therein result in weak non-specific binding only (Figure 1C). The same result is obtained when this motif is mutated in the context of a RAD51 promoter fragment. The latter is also a confirmed SOG1 target and contains a binding motif complying to the Bourbousse motifs including the CTT(N)_7AA_G sequence (Figure 1D, Supplementary Figure S1, Table 1). The loss of specific DNA binding likely explains why the same mutations in the *RAD51* promoter *in planta* result in the ΔRAD51 phenotype of *A. thaliana* transgenic plants upon genotoxic stress (19).

### SOG1^NAC^ binds to *pBCRA1* as a dimer with nanomolar aUinity

Generally, NAC transcription factors form homodimers and bind to DNA as a dimer. Using analytical SEC, the apparent molecular weight of the SOG^NAC^ in solution was determined to be 55.7 kDa (Supplementary Figure S3A). This value is larger than the theoretical value of 39.4 kDa for a homodimer and may result from the non-globular shape of the SOG1^NAC^ dimer. To assess the stoichiometry and affinity of SOG1^NAC^ interacting with DNA, ITC was performed with the 31 bp *pBRCA1* ds oligo containing the binding site and the non-target dsDNA fragment of the same length (Figure 2A&B). The ITC results confirm the specificity of SOG1^NAC^ for the *pBRCA1* DNA observed in EMSA experiments. Furthermore, the results show that the binding of the NAC domain to the DNA is an exothermic reaction and that SOG1^NAC^ binds as a dimer to the DNA, which is in line with the behaviour of other NAC TFs (33, 34). The dissociation constant (K_D)_ was determined to be 145 nM and the reaction is enthalpy driven (Figure 2A&C). AlphaFold3 structure predictions of SOG1^NAC^ bound to *pBRCA1* DNA suggests a cluster of positive charges on the NAC surface to be involved in DNA binding (Figure 2D), similar to what was previously observed for ORE1 (34). These positively charged regions are presumed to give rise to the non-specific, transient DNA binding observed in the EMSA experiments. In consequence, altering the buffer salt content and thus the ionic strength influences the affinity of the interaction between SOG1^NAC^ and *pBRCA1* (Supplementary Figure S4).

### SOG1^NAC^ specifically targets the Bourbousse binding motifs

The CTT(N)_7AA_G Ogita motif covers only half of the *in vivo* identified targets of SOG1, suggesting that the presence of this motif alone does not suffice as sole SOG1 binding prerequisite. In contrast, the Bourbousse motifs are more elaborate.

The *pBRCA1* site that was extensively used in this research thus far, matches the Ogita motif as well as all three Bourbousse motifs. The promoter region of the *SMR5* gene (*pSMR5*), another *in vivo* target of SOG1 (19, 35) contains three potential SOG1 binding sites (Table 1). Of these, site 1 contains the CTT(N)_7AA_G Ogita motif and is also compatible with the Bourbousse motifs. Site 2 does not contain CTT(N)_7AA_G, but is still compatible with a Bourbousse motif. Finally, site 3 does again contain the CTT(N)_7AA_G Ogita motif, but is otherwise not compatible with any of the Bourbousse motifs (Supplementary Figure S1). An EMSA with *pSMR5* dsDNA shows two bands of protein-DNA complex, suggesting the presence of only two functional binding sites (Figure 3A).

**Figure 3:**
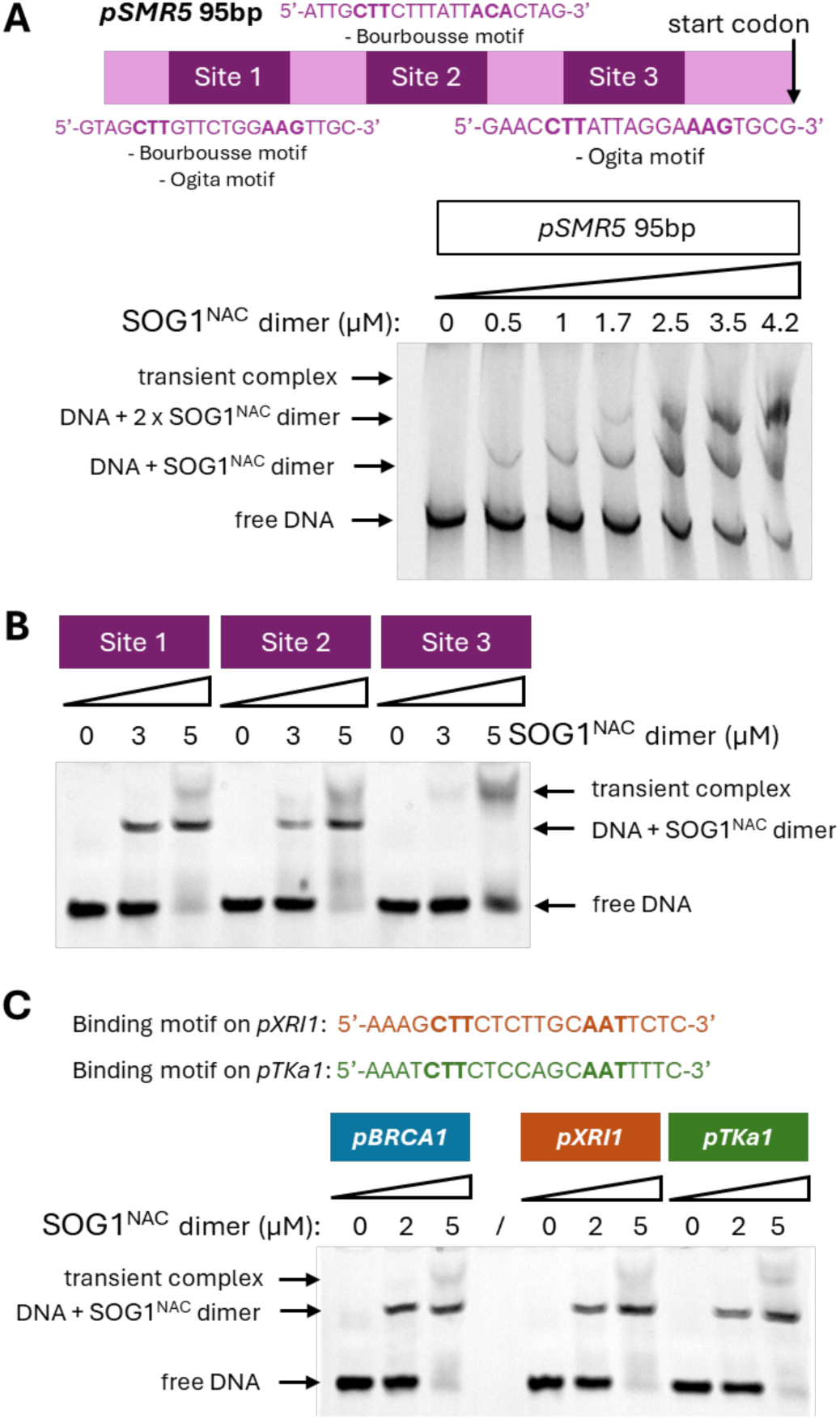
The Bourbousse identified motifs best describe the SOG1^NAC^ binding site. (A) Putative binding motifs present on pSMR5: site 1 and 2 contain Bourbousse motifs, while site 1 also confirms to the Ogita motif. Site 3 only contains an Ogita motif. EMSA of SOG1^NAC^ with a 95 bp dsDNA fragment of pSMR5 containing all three sites. (B) EMSA of SOG1^NAC^ with 31 bp dsDNA fragments of the three putative binding sites present on pSMR5, showing only aZinity for the Bourbousse motifs. (C) Binding motifs present on pXRI1 and pTKa1, both containing a Bourbousse motif without including an Ogita motif. EMSA of SOG1^NAC^ with 31 bp dsDNA fragments of pXRI1 and pTKa1.

To detect to which two sites SOG1^NAC^ binds, EMSAs were performed with shorter dsDNA fragments that each contain one of the three putative binding sites (Figure 3B). These show that SOG1^NAC^ only binds to the two sites that contain an identified Bourbousse motif and that the CTT(N)_7AA_G Ogita motif appears to be too exclusive to define the binding site as no specific binding is observed.

Further confirmation of the Bourbousse motifs as the true interaction sites for SOG1^NAC^ comes from EMSA experiments utilizing *XR1* and *TKA1* promoter fragments (*pXRI1* and *pTKa1*). These *in vivo* SOG1 targets both encompass Bourbousse motifs (Supplementary Figure S1) without including the CTT(N)_7AA_G motif (G to T substitution in final position of the motif; Figure 3C) (19). SOG1^NAC^ is able to bind both ds promoter fragments with a similar affinity as for the *pBRCA1* binding site. This confirms that the more general motifs defined by Bourbousse and co-workers better describe the target binding site of SOG1 and that the strict CTT(N)_7AA_G consensus site is too restrictive. Additional base pairs surrounding the CTT(N)_7AA_G motif likely also play a role and variations on this motif are also possible within this context.

### SOG1^NAC^ undergoes nucleic acids-driven phase separation *in vitro*

To better understand our observation that mixtures of purified SOG1^NAC^ and nucleic acids become turbid over time, the optical density at 390nm (OD_390)_ was measured after mixing SOG1^NAC^ with poly-A RNA. SOG1^NAC^ evolves a turbidity profile that is typical for many phase separating proteins: an increase of OD_390 fo_llowed by a decrease that stabilizes to a baseline (Figure 4A) (36). From this data, a re-entrant phase behaviour is observed. The OD_390nm ma_xima that are reached in the OD_390 ve_rsus time profiles increase with rising concentrations of poly-A, but decline starting at concentrations poly-A between 100-250μg/ml until the signal eventually no longer detectable is (Figure 4B). Formation of spherical droplets for varying protein and poly-A concentrations was confirmed using fluorescence microscopy (Figure 4C and Supplementary Figure S5). Through dual colour labelling, SOG1^NAC^ and poly-A were demonstrated to co-integrate into the droplets (Figure 4D). Interestingly, both droplets with a hollow shell-like fluorescence pattern and droplets with an even solid fluorescence pattern are observed (Figure 4E). Variation of the ionic strength does influence turbidity profiles of SOG1^NAC^-poly-A mixtures indicating that electrostatic interactions are involved in the phase behaviour (Figure 4F), as was also seen for the DNA binding experiments. In contrast to salt concentrations between 68 and 100 mM NaCl, a substantial drop in turbidity is observed for higher salt concentrations containing 150 mM NaCl. Droplet growth in function of time was mapped by DLS for 4 hours (Figure 4G). The hydrodynamic radius R_h in_ function of time follows Ostwald ripening for the first two hours (Kraska 2008), highlighting the dynamic and liquid-like nature of the droplets that are initially formed (37). After two hours, the droplets size stabilizes. FRAP of a ten minute old reaction shows limited recovery of 27 percent, two minutes post bleaching (Figure 4H). Remark that for a subset of droplets recovery was observed exclusively in their outer layers, reminiscent of the previously mentioned droplets with shell-like fluorescence pattern. This indicates that while there is a significant mobile fraction present in the condensates, there is also a less mobile fraction present.

**Figure 4:**
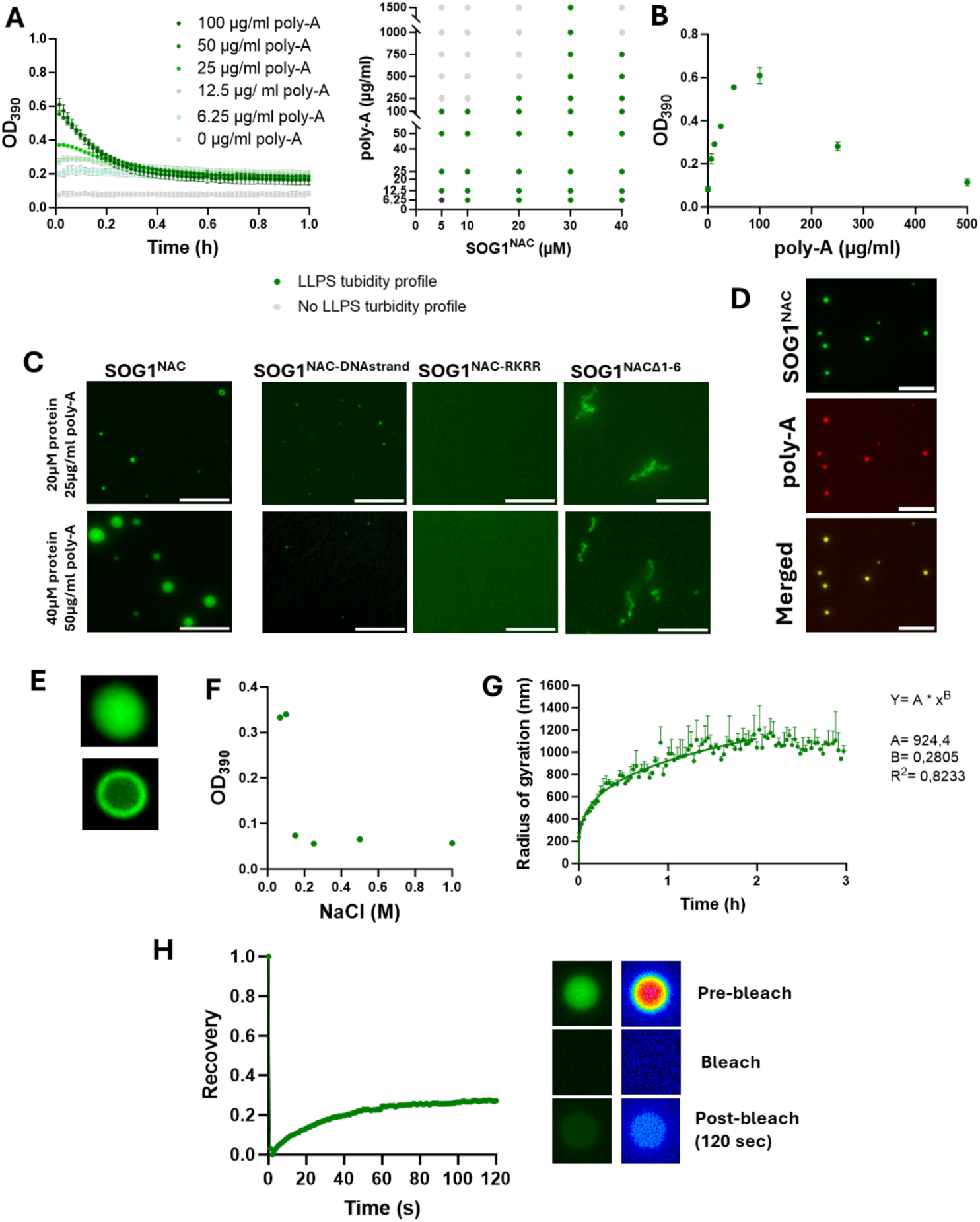
RNA-dependent phase behaviour of SOG1^NAC^. (A) Turbidity profiles of NAC in function of time at 390nm (left panel) and phase-diagram of NAC at diZerent concentrations poly-A based on turbidity profiles (right panel). (B) OD390 versus poly-A concentrations: re-entrant phase behaviour with increasing OD390 signals for increasing poly-A concentrations that drop again for even higher concentrations. (C) Fluorescence microscopy image of DyLight488-labeled protein (SOG1^NAC^, SOG1^NAC –DNAstrand^ mutant, SOG1^NAC - RKRR^ mutant and SOG1^NACΔ1-6^) in presence of poly-A RNA for diZerent concentrations. (D) Fluorescence microscopy image of co-integration of DyLight488-labeled NAC and Cy5-labeled poly-A RNA in spherical droplets. (E) shell-like and fully coloured droplets of DyLight488 labelled NAC in presence of unlabelled poly-A. (F) OD390 versus NaCl concentrations for reactions with 20 µM SOG1^NAC^and 25 µg/ml poly-A. (G) Droplet growth of a reaction with 20 µM SOG1^NAC^ and 25 µg/ml poly-A followed by dynamic light scattering. (H) FRAP of a droplet with 20 µM SOG1^NAC^ and 25 µg/ml poly-A (green fluorescence on the right and heat map on the left).

Next, we investigated if addition of dsDNA to SOG1^NAC^ similarly could induce phase separation. DNA fragments used include the full 310 bp promoter sequence of BRCA1, the 31 bp *pBRCA1* ds oligo containing the Ogita/Bourbousse binding motif and the 293 bp promoter of the non-target house-keeping gene UBQ10 (Figure 5A). Small condensates were formed in SOG1^NAC^ mixtures with the short 31 bp *pBRCA1* fragment, while shell-like droplets and irregular structures (called clusters) are formed in presence of the longer 310 bp fragment or the 293 bp non-target sequence. The higher the concentration of dsDNA, the more cluster-like structures are present (Figure 5B). Interestingly, FRAP experiments confirm a dynamic behaviour of those droplet clusters with a recovery of around 80 % after two minutes (Figure 5C). This discrepancy in recovery rate between dsDNA and poly-A driven LLPS may indicate a different internal structure.

**Figure 5:**
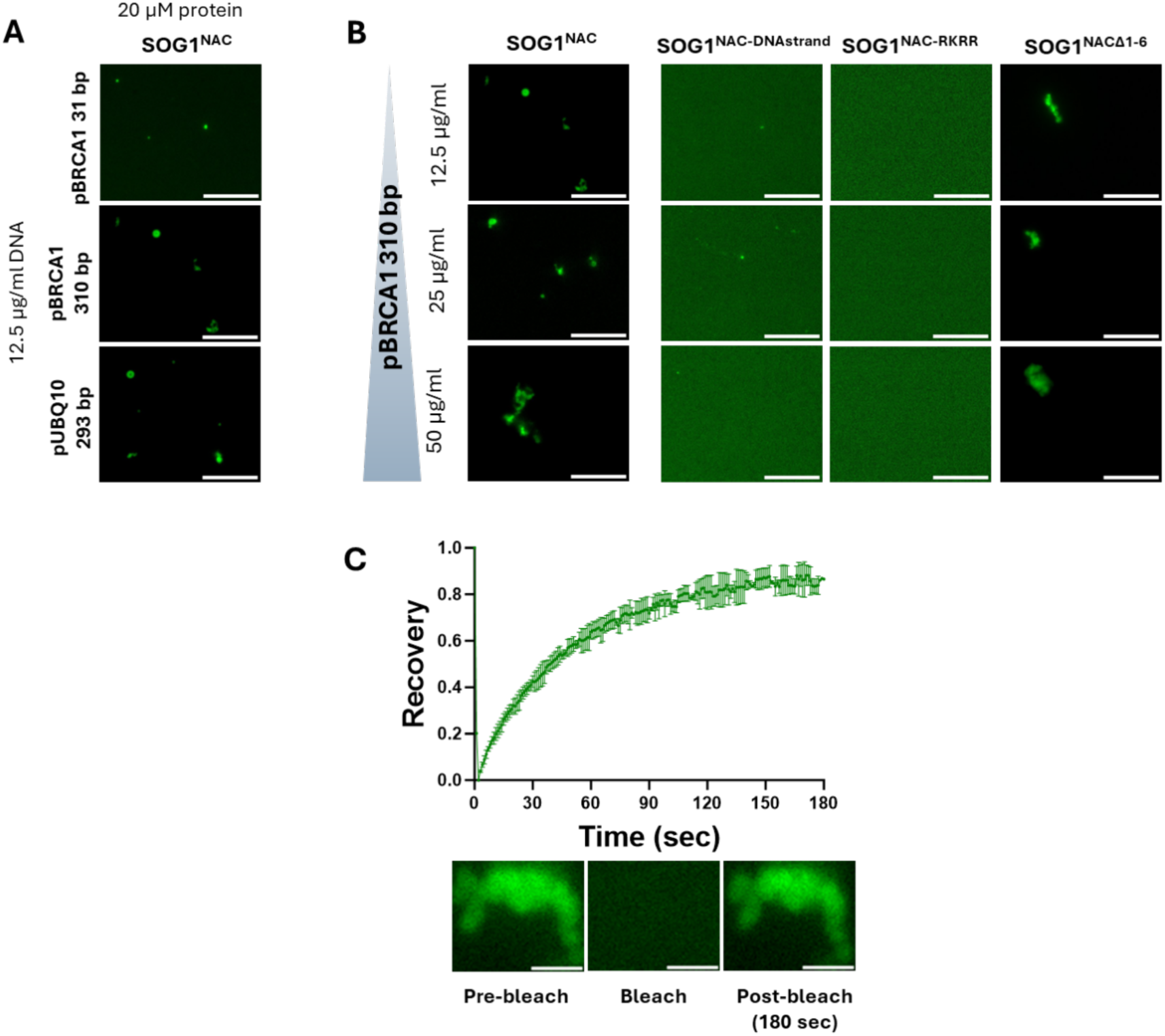
DNA-driven phase separation of SOG1^NAC^ and its mutants (A) with pBRCA1 31 bp oligo, and target versus non-target DNA promoter sequences. (B) with diZerent concentrations of the promoter sequence of BRCA1 target. (C) FRAP on irregular clusters of 20µM NAC and 12.5 µg/ml BRCA1 promoter sequence (310 bp).

### Dimerization of the SOG1 NAC domain is required for spherical droplet formation and proper DNA-binding

We next investigated whether dimerization is required for SOG1^NAC^ DNA-binding and phase separating behaviour. Based on AlphaFold3 structural predictions of the SOG1^NAC^ dimer, a truncation mutant lacking the N-terminal β-strand (ΔL58-K63) was designed (from now called SOG1^NACΔ1-6^). The SOG1^NACΔ1-6^ protein has an estimated molecular weight of 19.8 kDa as measured using analytical SEC, confirming a (globular) monomeric state in contrast to the non-truncated SOG1^NAC^ that eluted as a dimer (Supplementary Figure S3A). The CD-profiles of SOG1^NAC^ and SOG1^NACΔ1-6^ are similar with a negative peak around 220 nm indicating the presence of predominantly β-sheets in both proteins (Supplementary Figure S3B).

Despite being monomeric, SOG1^NACΔ1-6^ still binds to the 31 bp *pBRCA1* ds oligo, but with lower affinity compared to wild-type SOG1^NAC^ (Figure 6A&B). The affinity and stoichiometry could not be further assessed by ITC, because the SOG1^NACΔ1-6^ protein is too unstable at the higher concentrations used for this. This implies that dimerization is needed for the intrinsic stability of the isolated NAC domain. The decrease in affinity is likely due to the independent binding of the SOG1^NACΔ1-6^ monomers compared to the avidity generated via a stable dimer, although a lower thermodynamic stability of SOG1^NACΔ1-6^ may also play a role.

**Figure 6:**
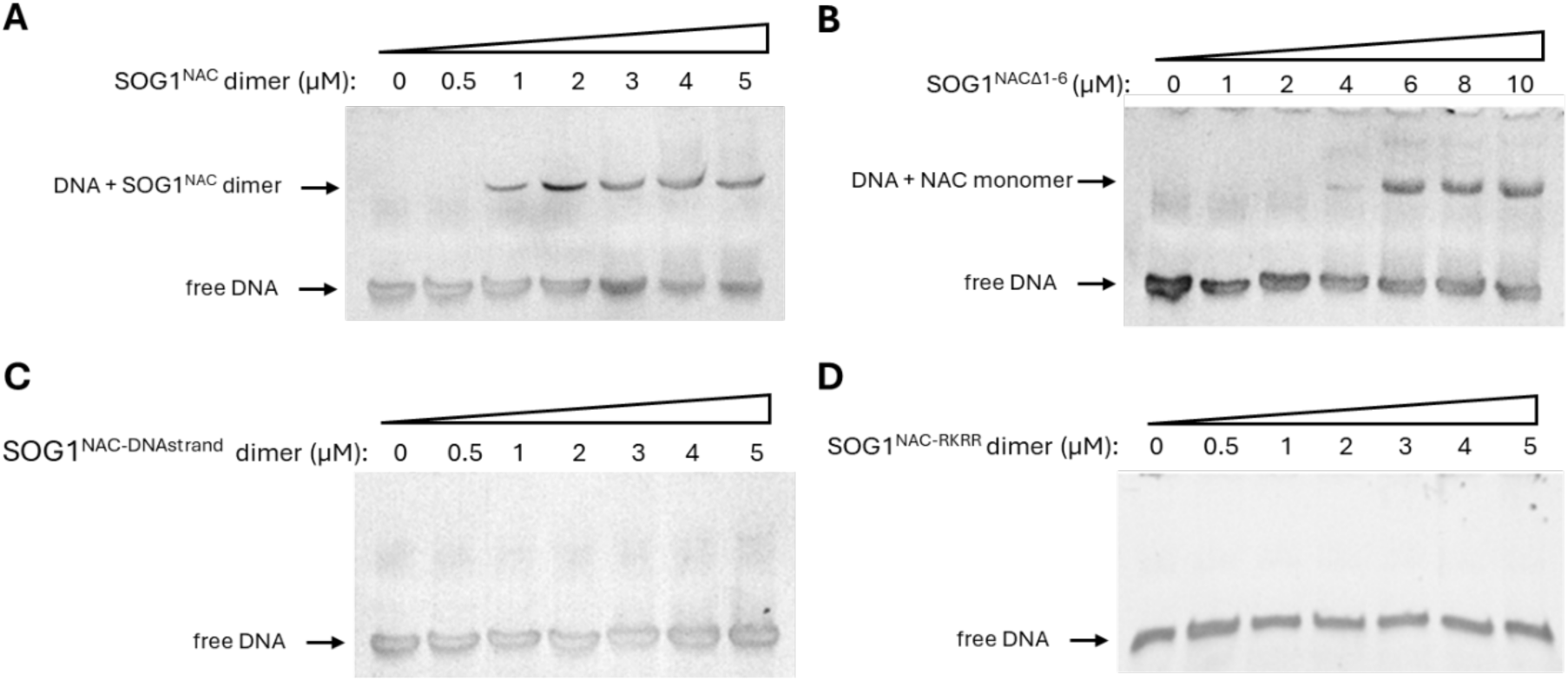
DNA binding of SOG1^NAC^ and its mutants. (A), (B), (C) & (D) EMSA of WT SOG1^NAC^, SOG1^NACΔ1-6^ (=monomer), the SOG1^NAC-DNAstrand^ mutant and the SOG1^NAC-RKRR^ mutant, respectively, with a 31 bp ds oligo of pBRCA1 containing the SOG1^NAC^ binding site.

SOG1^NACΔ1-6^ shows a changed LLPS behaviour as the formation of spherical droplets in presence of poly-A is no longer observed. Instead, irregular structures are formed (Figure 4C and Supplementary Figure S5). In presence of DNA, phase separation is again defective. No spherical droplets are observed, but occasionally cluster-like structures appear. This suggest that dimerization is required for spherical droplet formation by both RNA- or and DNA-driven LLPS.

### Two positively charged clusters of SOG1^NAC^ link DNA-binding with RNA- and DNA-driven LLPS

AlphaFold3 structure predictions of SOG1^NAC^ bound to the 31 bp *pBRCA1* fragment reveal two segments rich in positively charged amino acids that likely determine DNA specificity due to their proximity to the nucleotides making up the target binding site. These are a loop encompassing residues R136-R139 and a β-strand encompassing residues R150-T157 (residues positioned on the full length protein) (Supplementary Figure S6). The corresponding segments in the NAC domains of ANAC019 and ORE-1 differ in amino acid sequence, but also constitute the DNA binding site (33, 34). Two SOG1^NAC^ mutants harbouring mutations in these two putative DNA binding segments were designed. For both mutant proteins, the positive charges were mutated to non-charged amino acids. The first mutant is referred to in this paper, is the SOG1^NAC-RKRR^ mutant that has following mutations: R79A, K80A, R81A, and R82A. The second is called the SOG1^NAC-DNAstrand^ mutant with mutations R93S, H95S, T97A, R99S, and T100S, which are located in the β-strand that docks into the major DNA-groove. CD spectra of both mutants are very similar to that of the WT SOG1^NAC^ protein, indicating proper folding (Supplementary Figure S3B). Both elute at approximately the same volume as wild-type SOG1^NAC^ in analytical SEC: apparent molecular weights of 67, 4 and 58, 2k Da for SOG1^NAC-RKRR^ and SOG1^NAC-DNAstrand^ mutant, respectively. We therefore assume that, like SOG1^NAC^, the mutants maintain their ability to form homodimers in solutions (Supplementary Figure S3A).

EMSA experiments performed with both mutants and the 31 bp *pBRCA1* fragment show a complete loss of DNA binding capabilities (Figure 6C&D). This indicates that both structural elements that were mutated in these NAC domain variants are essential parts of the DNA binding site. Furthermore, it implies that SOG1^NAC^ employs a similar binding mechanism as ANAC019 and ORE-1 that is presumably conserved within the entire family of NAC transcription factors (33, 34).

We next investigated if the cluster of positive charges that is essential for DNA binding is also DNA- or RNA-mediated phase separation (Figure 5B). DNA-mediated phase separation is abolished for both mutants, confirming that an intact DNA binding site, or at least the presence of positive charges, is essential for multivalent protein-DNA interactions.

RNA-driven phase behaviour of the two mutants is altered compared to wild-type SOG1^NAC^. However, RNA-induced phase behaviour is not completely abolished. For the SOG1^NAC-DNAstrand^ mutant, small droplets are still formed in most of the conditions tested (Figure 4C and Supplementary Figure S4). The intact RKRR loop is suggested to on its own be able to support the LLPS potential of the SOG1^NAC-DNAstrand^ mutant with DNA. Altogether, these results establish a functional link between the DNA-binding potential of SOG1^NAC^ and its ability to undergo phase separation both with RNA and DNA.

### The phase-separating behaviour of SOG1^NAC^ is controlled by a tight interplay between DNA and RNA binding

The results above suggest that both RNA and DNA bind to overlapping regions on the surface of SOG1^NAC^. Therefore, the question follows whether DNA and RNA can co-integrate in the SOG1^NAC^ condensates or if one would exclude the other. We first challenged poly-A-SOG1^NAC^ condensates with the addition of *pBRCA1* 31 bp oligo. Below equimolar concentrations SOG1^NAC^ dimer and DNA, the DNA fragment can integrate in the RNA-induced condensates (Figure 7A). However, increasing *pBRCA1* 31 bp oligo concentrations weakens the RNA-driven phase behaviour. At equimolar concentrations of SOG1^NAC^ dimer and the *pBRCA1* 31 bp target sequence, poly-A induced condensates of SOG1^NAC^ are fully dissolved. The pBRCA1 31 bp DNA target competes with RNA and prevents poly-A driven condensation by excluding poly-A from the droplets. This agrees with the hypothesis that poly-A and the *pBRCA1* 31 bp oligo bind to the same or overlapping sites on SOG1^NAC^. Binding of the *pBRCA1* 31 bp oligo to SOG1^NAC^ presumably breaks the multivalent dynamic SOG1^NAC^-RNA interactions, that stabilize the condensates, although few small condensates still form.

**Figure 7:**
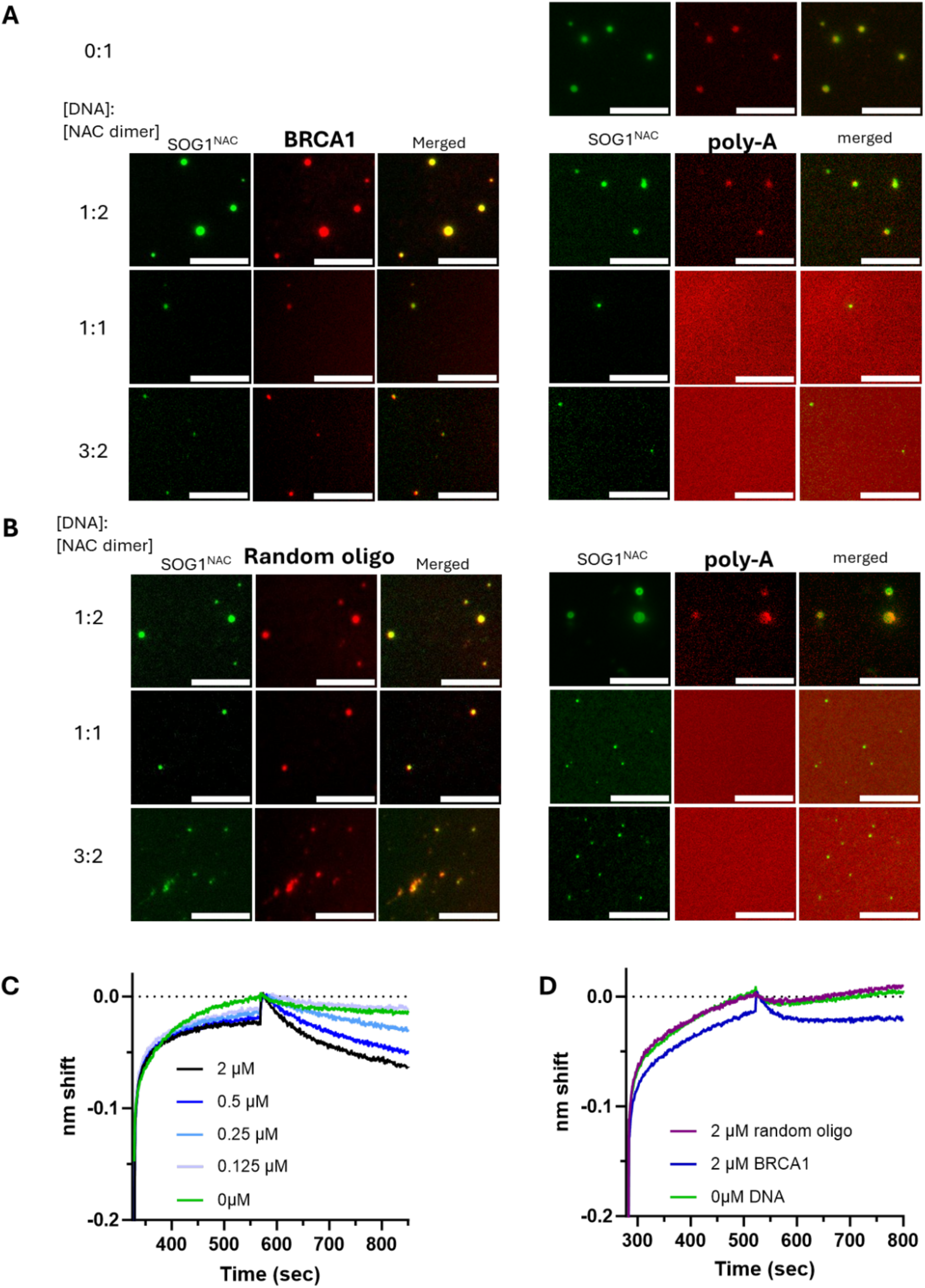
Competition assays RNA versus DNA-binding of SOG1^NAC^ - Fluorescence microscopy to visualize RNA and DNA cointegration in poly-A-induced SOG1^NAC^ condensates. 10 µM SOG1^NAC^dimer (green fluorescence) in presence of 25 ug/ml poly-A (left: unlabelled, right: red fluorescence) was titrated with various amount DNA (left: red fluorescence, right: unlabelled), either pBRCA1 31 bp DNA oligo (A) or Random 31 bp DNA oligo (B). Bio-layer interferometry of immobilized SOG1^NAC^ and poly-A RNA versus various concentrations pBRCA1 31 bp DNA oligo (C) and random non-target DNA oligo compared to pBRCA1 31 bp DNA oligo and no DNA (D).

When this competition experiment is repeated with a non-specific DNA fragment of the same length (Figure 7B), again poly-A is excluded from the droplets at a 1:1 molar ratio. Differently, more droplets are assembled with SOG1^NAC^ and non-target DNA at this equimolar ratio. Probably, no stable soluble SOG1^NAC^-non-cognate DNA complexes are formed and transient and weak multivalent protein-DNA interactions involving a non-cognate DNA sequence enable DNA-driven condensation.

The finding that DNA with a specific SOG1-binding motif can outcompete RNA is supported by a biolayer-interferometry (BLI) competition assay (Figure 7C). After association of poly-A to immobilized SOG1^NAC^, subsequent dipping in *pBRCA1* 31 bp oligo results in a dissociation profile for which the observed k_o? ra_te is clearly higher than when poly-A is released from SOG1^NAC^ in absence of the DNA fragment. For non-target DNA, no such dissociation is observed (Figure 7D), thus in line with ITC results where no binding is detected for non-target DNA.

## Discussion

SOG1 is a plant transcription factor central to the DNA damage response in which it takes on the role of mammalian p53 (23). Upon genotoxic stress, it controls the transcription of around 300 genes, either directly or indirectly. The target binding sites of SOG1 have been previously studied *in vivo* using ChIP-sequencing and RNA sequencing by two groups independently resulting in somewhat similar conclusions (19, 20). Here we show that SOG1^NAC^ target sequences are best described by the Bourbousse DNA motifs enriched in the gene groups that are mostly upregulated by SOG1 upon genotoxic stress. The CTT(N)_7AA_G motif identified by Ogita et al., 2018 is too restrictive as this motif can be perturbed by two nucleotides without loss of SOG1^NAC^ binding and residues outside of the CTT(N)_7AA_G motif also affect *in vitro* binding.

As observed for other NAC TFs, SOG1^NAC^ binds to DNA as a dimer. Truncated, monomeric SOG1^NAC^ binds target DNA with substantially decreased affinity. This is likely due to the avidity advantage of the dimer or loss of intrinsic stability of the monomer. In addition, mutating two positively charged segments that are implicated in DNA recognition in ANA019 and ORE1 (RKRR and DNA-strand regions on SOG1^NAC^) abolish DNA binding. From this we concluded that SOG1^NAC^ uses a similar mechanism for operator recognition as other NAC transcription factors which is canonical for this family (33, 34).

We further show that SOG1^NAC^ displays phase separation *in vitro* in presence of RNA as well as DNA. Of interest here is that SOG1^NAC^ is a fully folded, globular protein. The majority of the available literature as of yet typically focuses on the importance of intrinsically disordered and low complexity domains in LLPS, downplaying the contribution of folded domains within a larger LLPS-prone protein containing IDR regions (for a review see (16)). Functional LLPS of fully folded proteins is rare. While LLPS can typically be induced at high concentrations using crowding agents as initially observed for lysozyme ((38)), only few do so under physiological conditions, a rare example being *E. coli* Lipoate-protein ligase A (39). Folded domains typically drive LLPS when present as repeats in a beads-on-a-string format that creates multivalency or when interactions with LLPS partners induce unfolding (for a review see: (16)).

Phase separation by SOG1^NAC^ involves a fully folded dimer. The potential of SOG1^NAC^ to phase separate in presence of nucleic acids is shown to be functionally linked to its ability for specific DNA binding and dimerization, and therefore seems to require a folded state.

Monomeric SOG1^NAC^ does not form spherical condensates, and dimerization therefore contributes to the multivalency required to phase separate. Dimerization via a folded domain has been shown to be essential to enable LLPS of other proteins such as G3BP1 in the RNA-driven assembly of stress granules (40). Dimerization by its NTF2L domain allows to increase the valency of the separate RNA-binding domain. For SOG1^NAC^ both functions are sequestered on a single folded entity.

LLPS is a key component of regulation of transcription and translation, involving a complex interplay between the components involved: transcription factors, DNA, RNA polymerase, RNA, the mediator complex, chromatin and a variety of other regulatory proteins (7, 41). The RNA-dependent phase behaviour of the conserved SOG1^NAC^ implies an RNA-mediated feedback control mechanism for the formation of condensates at transcription start sites, corresponding to the model proposed by Henninger *et al.* (2021) (11). According to this model, condensate formation at transcription sites is abolished in the presence of too high concentrations of mRNA. The latter is mimicked by the re-entrant phase behaviour of SOG1^NAC^seen in presence of poly-A. A similar re-entrant RNA-dependent phase behaviour was observed for the human androgen receptor where its folded DNA-binding domain serves as the minimal region driving LLPS (14).

A tight and complex interplay between DNA (target versus non-target) and RNA binding controls the formation of condensates at transcription sites. For SOG1^NAC^, RNA and dsDNA compete with each other for LLPS. Mutations that abolish specific DNA binding of SOG1^NAC^ also induce an altered DNA- and RNA-dependent phase separation behaviour compared to the wild-type protein. This suggests that the binding site for specific DNA binding overlaps with that for both non-specific DNA and RNA recognition that drives LLPS, allowing DNA specificity to influence the LLPS behaviour of SOG1^NAC^. This agrees with similar differential behaviour of other transcription factors where both RNA- and DNA-mediated condensation benefit from sequence-specific interactions (for a review see (42)).

The interplay between RNA- and DNA-induced LLPS is an essential part of transcription; typically transcription factors first undergo DNA-driven LLPS to initiate transcription, while the resulting RNA accumulation subsequently transit the TFs into RNA-driven LLPS. The behaviour of SOG1 with respect to the competition between RNA and DNA reported on in this manuscript reflects this switch in LLPS driving force. Competition of DNA and RNA for a shared or overlapping binding site may form the structural basis for the transition between two types of transcription-related condensates.

## Acknowledgments

We thank Dr. Indra Bervoets for the initiation of the radioactive EMSA experiments and Sarah Haesaerts for helping with the purification of the proteins.

