## Supplemental table and figures for "DNA-binding and dimerization of the SOG1 NAC domain are functionally linked with its ability to undergo liquid-liquid phase separation"

- Supplementary Figure S1: SOG1 binding motifs identified by Bourbousse *et al.*
- Supplementary Figure S2: $SOG1^{NAC}$ targets the binding site located on *pBRCA1*
- Supplementary Figure S3: Dimerization and secondary structure composition of $SOG1^{NAC}$ and its mutants
- Supplementary Figure S4: Ionic strength dependence of the interaction between $SOG1^{NAC}$ and target DNA
- Supplementary Figure S5: Fluorescence microscopy image of DyLight488-labeled proteins
- Supplementary Figure S6: $SOG1^{NAC}$ in complex with DNA

**Supplementary Table legends**

- Supplementary Table 1: Primers for mutagenesis of $SOG1^{NAC}$ clone

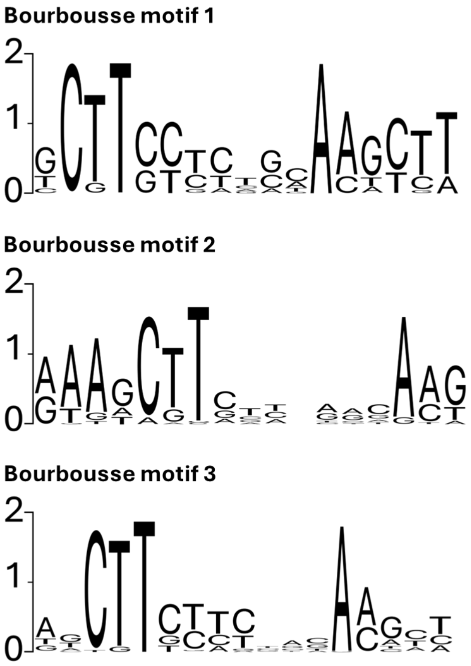


**Supplementary Figure S1: SOG1 binding motifs identified by Bourbousse et al..** Motifs 1, 2 and 3 represent the enriched sequence motifs identified on the promoters of the gene groups that are mostly, second mostly and third mostly upregulated by SOG1 upon genotoxic stress, respectively. Figures are based on data taken from Bourbousse et al., 2018.

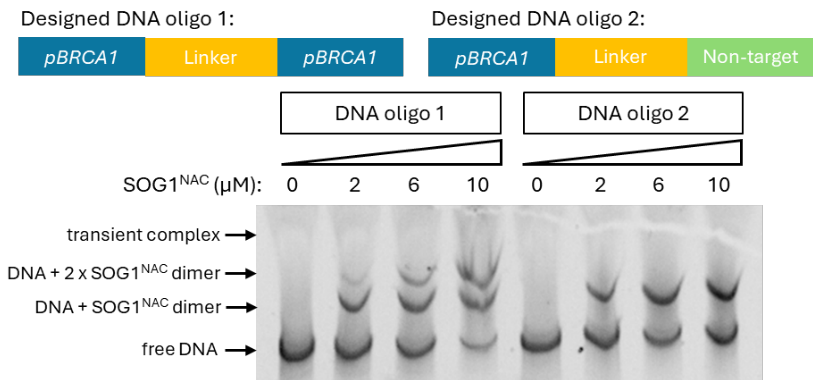


**Supplementary Figure S2: SOG1$^{NAC}$ targets the binding site located on pBRCA1.** EMSA of SOG1$^{NAC}$ with two designed DNA oligos. DNA oligo 1 contains 31 bp oligos of the pBRCA1 binding site connected by a random linker. DNA oligo 2 contains the same 31 bp pBRCA1 fragment connected by the same linker to a 31 bp randomly generated, non-target DNA fragment.

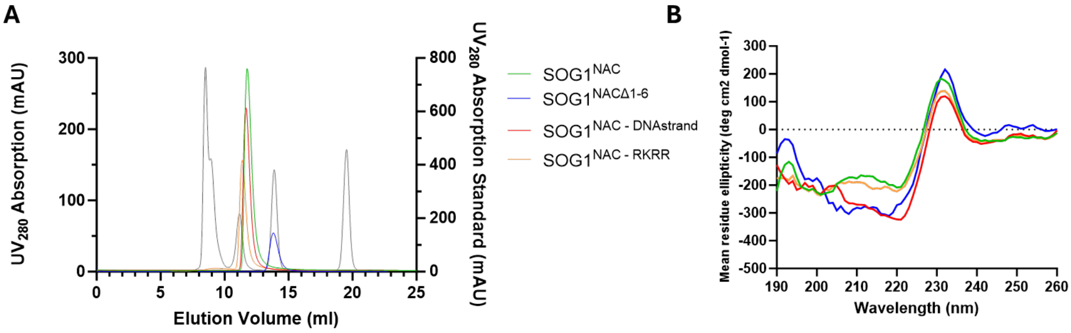


**Supplementary Figure S3: Dimerization and secondary structure composition of SOG1$^{NAC}$ and its mutants.** (A) Analytical SEC of SOG1$^{NAC}$ and its mutants. Graph shows the elution volumes of each protein compared to the elution volumes of the molecular weight standard proteins (grey). (B) CD spectra of SOG1$^{NAC}$ and its mutants, showing similar profiles. The positive peaks at 230 nm may be linked to aromatic-aromatic interactions or disulfide bounds (43).

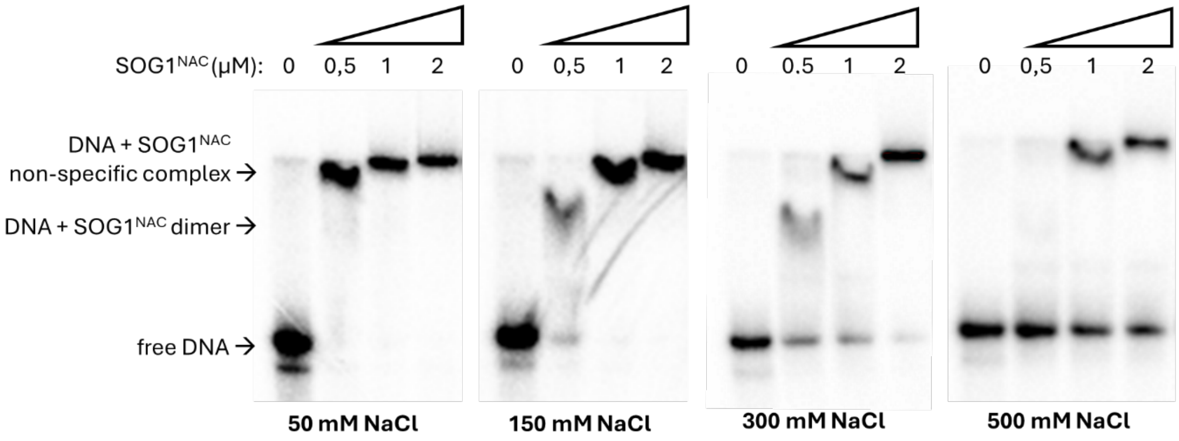


**Supplementary Figure S4: Ionic strength dependence of the interaction between SOG1$^{NAC}$ and target DNA.** Radioactive EMSAs of SOG1$^{NAC}$ with 99 bp pBRCA1 DNA using different salt concentrations. From left to right: EMSA reactions with 50, 150, 300 and 500 mM NaCl containing binding buffers.

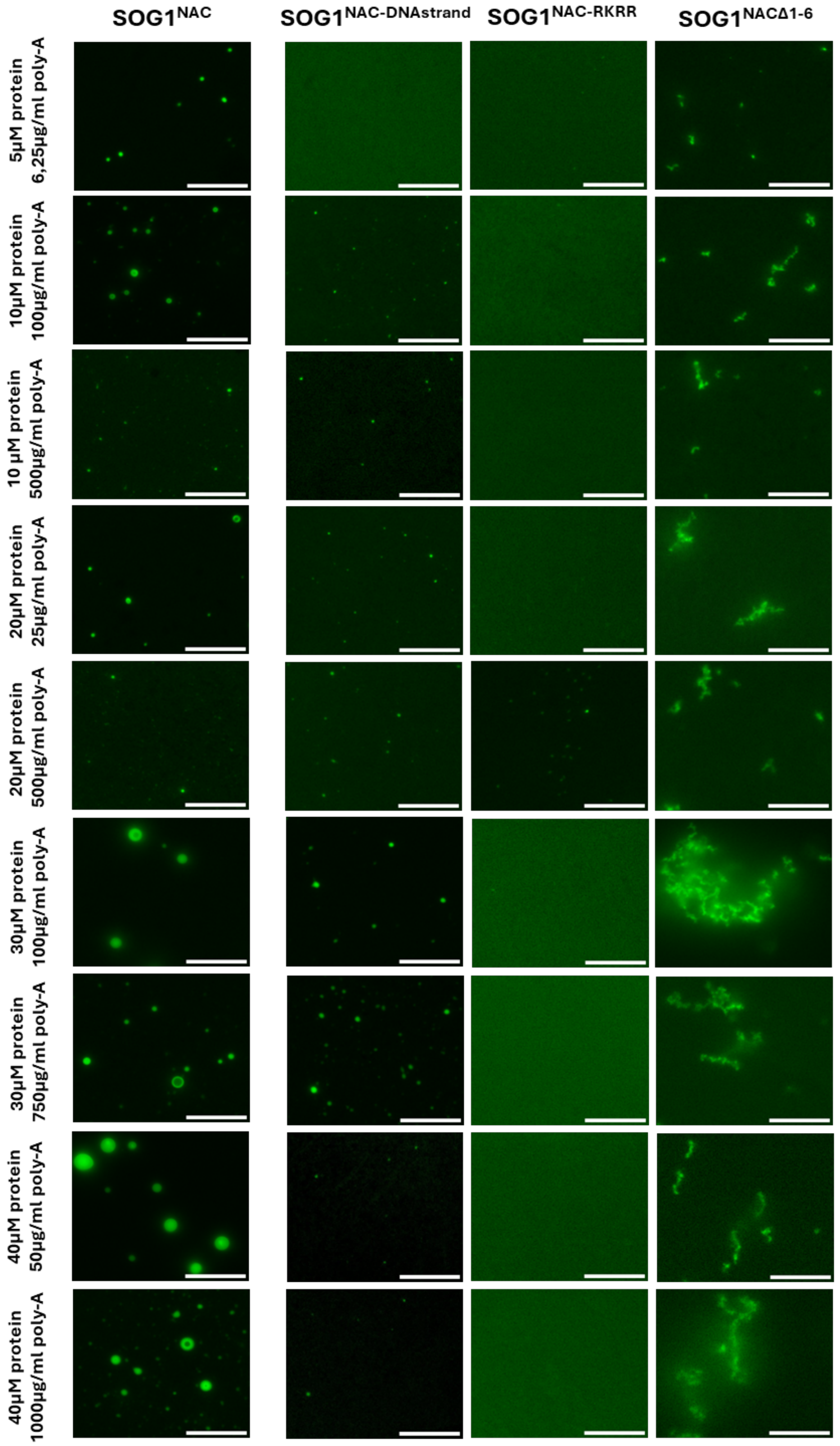


**Supplementary Figure S5: Fluorescence microscopy image of DyLight488-labeled protein.** Shown are NAC, SOG1$^{NAC\text{-}DNAstrand}$ mutant, SOG1$^{NAC\text{ - }RKRR}$ mutant and SOG1$^{NAC\Delta1\text{-}6}$ in presence of different concentrations of poly-A RNA.

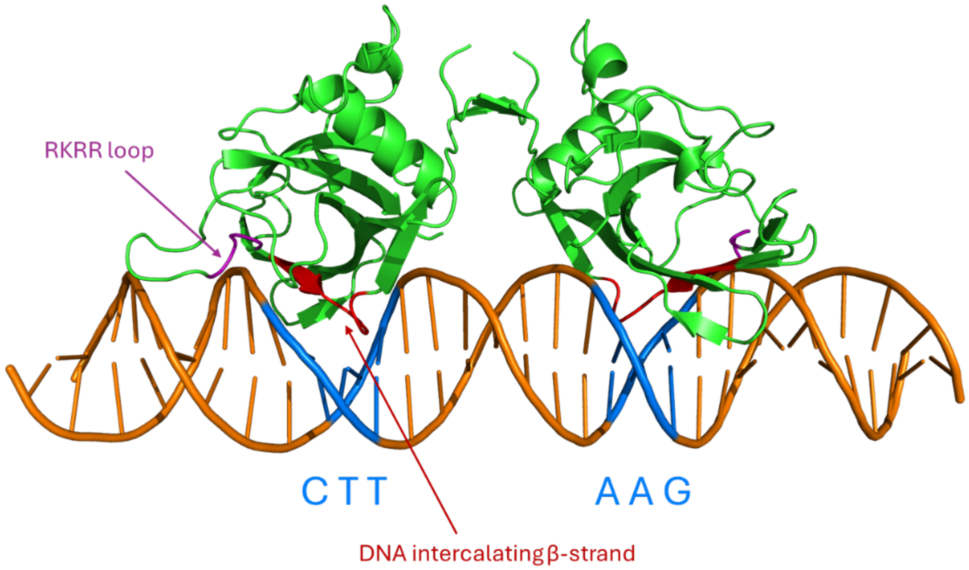


**Supplementary Figure S6: SOG1$^{NAC}$ in complex with DNA.** AlphaFold3 structure prediction of SOG1$^{NAC}$ binding to a 31 bp fragment of pBRCA1. The six CTT-AAG nucleotides that are separated by the $N_7$-linker in the Ogita CTT(N)$_7$AAG motif are marked in blue. The RKRR-loop that is in close proximity to the DNA and the β-strand that intercalates with the DNA are marked in purple and red, respectively. The AAs of both structural elements are mutated to create the SOG1$^{NAC}$, SOG1$^{NAC-RKRR}$ and SOG1$^{NAC-DNAstrand}$ mutants.

**Supplementary Table S1: Primers used for mutagenesis of SOG1$^{\text{NAC}}$**

| Primer name | Sequence (5'- … -3') |
|---|---|
| *XbaI FW* | GAGCGGATAACAATTCCCCTCTAGAAATAATTTTGTTTAACTTT |
| SOG1$^{\text{NAC}\Delta\text{1-6}}$ *FW* | TGTATTTCCAAGGCCGTGGCTTTGACCCGAGCGACCCGGAGATTATC |
| SOG1$^{\text{NAC}\Delta\text{1-6}}$ *RV* | ATCTCCGGGTCGCTCGGGTCAAAGCCACGGCCTTGGAAATACAGGTTC |
| SOG1$^{\text{NAC-RKRR}}$ FW | AAGCGTATAGCACCGGCACCGCTGCCGCAGCTAAAATTCACGACGATGACTTCGGTGA |
| *SOG1$^{NAC\text{-}RKRR}$ RV* | AAGTCATCGTCGTGAATTTTAGCTGCGGCAGCGGTGCCGGTGCTATACGCTTTGATC |
| *SOG1$^{NAC\text{-}DNAstrand}$ FW* | ACGATGACTTCGGTGATGTTTCTTGGAGCAAGGCCGGCTCATCGAAACCGGTGGTTCTGGACGGTGTGCAG |
| *SOG1$^{NAC\text{-}DNAstrand}$ RV* | CCGTCCAGAACCACCGGTTTCGATGAGCCGGCCTTGCTCCAAGAAACATCACCGAAGTCATCGTCGTGA |
| *BamHI (to Acc651 mutated) RV* | TCGGGCTTTGTTAGCAGCCGGTACCTTACTGCTGATAGAAAATC |
